# Lipid associated polygenic enrichment in Alzheimer’s disease

**DOI:** 10.1101/383844

**Authors:** Iris J. Broce, Chin Hong Tan, Chun Chieh Fan, Aree Witoelar, Natalie Wen, Iris Jansen, Christopher P. Hess, William P. Dillon, Christine M. Glastonbury, Maria Glymour, Jennifer S. Yokoyama, Fanny M. Elahi, Gil D. Rabinovici, Bruce L. Miller, Elizabeth C. Mormino, Reisa A. Sperling, David A Bennett, Linda K. McEvoy, James B. Brewer, Howard A. Feldman, Danielle Posthuma, Bradley T. Hyman, Gerard S. Schellenberg, Kristine Yaffe, Ole A. Andreassen, Anders M. Dale, Leo P. Sugrue, Celeste M. Karch, Rahul S. Desikan

## Abstract

Cardiovascular (CV) and lifestyle associated risk factors (RFs) are increasingly recognized as important for Alzheimer’s disease (AD) pathogenesis. Beyond the ∊4 allele of apolipoprotein E (*APOE*), comparatively little is known about whether CV associated genes also increase risk for AD (genetic pleiotropy). Using large genome-wide association studies (GWASs) (total *n* > 500,000 cases and controls) and validated tools to quantify genetic pleiotropy, we systematically identified single nucleotide polymorphisms (SNPs) *jointly* associated with AD and one or more CV RFs, namely body mass index (BMI), type 2 diabetes (T2D), coronary artery disease (CAD), waist hip ratio (WHR), total cholesterol (TC), low-density (LDL) and high-density lipoprotein (HDL). In fold enrichment plots, we observed robust genetic enrichment in AD as a function of plasma lipids (TC, LDL, and HDL); we found minimal AD genetic enrichment conditional on BMI, T2D, CAD, and WHR. Beyond *APOE*, at conjunction FDR < 0.05 we identified 57 SNPs on 19 different chromosomes that were jointly associated with AD and CV outcomes including *APOA4, ABCA1, ABCG5, LIPG*, and *MTCH2/SPI1.* We found that common genetic variants influencing AD are associated with multiple CV RFs, at times with a different directionality of effect. Expression of these AD/CV pleiotropic genes was enriched for lipid metabolism processes, over-represented within astrocytes and vascular structures, highly co-expressed, and differentially altered within AD brains. Beyond *APOE*, we show that the polygenic component of AD is enriched for lipid associated RFs. Rather than a single causal link between genetic loci, RF and the outcome, we found that common genetic variants influencing AD are associated with multiple CV RFs. Our collective findings suggest that a network of genes involved in lipid biology also influence Alzheimer’s risk.

## INTRODUCTION

There is mounting evidence that cardiovascular (CV) disease impacts Alzheimer’s disease (AD) pathogenesis. Co-occurrence of CV and AD pathology is the most common cause of dementia among the elderly [4] and imaging manifestations of vascular disease are routinely observed on MRI scans of AD patients [36]. Observational epidemiology studies have found that cardiovascular/lifestyle related risk factors (RFs) are associated with dementia risk and targeting these modifiable RFs may represent a viable dementia prevention strategy [5, 26]. Recently, the National Academy of Medicine [24] and the Lancet [19] commissioned independent reports on strategies for dementia prevention. Both reports found encouraging evidence for targeting cardiovascular RFs with the Lancet commission concluding that 35% of dementia could be prevented by modifying several RFs including diabetes, hypertension, obesity, and physical inactivity.

Genetic studies have found CV associated loci that also increase risk for late-onset AD. The ∊4 allele of apolipoprotein E (*APOE*) is the biggest genetic risk factor for AD and encodes a lipid transport protein involved in cholesterol metabolism [22]. Genome-wide association studies (GWAS) in late-onset AD have identified single nucleotide polymorphisms (SNPs) implicated in lipid processes, such as *CLU* and *ABCA7*[17, 31], and enrichment in cholesterol metabolism pathways [7]. Considered together, these findings suggest ‘pleiotropy’, where variations in a single gene can affect multiple, seemingly unrelated phenotypes [37].

We have previously shown that genetic enrichment in CV traits/diseases (hereafter referred to as RFs) results in improved statistical power for discovery of novel AD genes [10]. Building on this work, in the present study, we systematically evaluated shared genetic risk between AD and cardiovascular/lifestyle associated RFs and diseases. We focused on publicly available genetic data from cardiovascular outcomes and a combination of traits and diseases that have been epidemiologically associated with increased AD risk. Using large GWAS and validated tools to estimate pleiotropy, we sought to identify SNPs *jointly* associated with AD and one or more CV RFs, namely body mass index (BMI), type 2 diabetes (T2D), coronary artery disease (CAD), waist hip ratio (WHR), total cholesterol (TC), low-density (LDL), and high-density lipoprotein (HDL). Using large publicly available databases, we additionally examined whether the AD/CV pleiotropic genes are enriched for biological processes, over-represented within specific tissue and cell types, co-expressed, and differentially expressed within AD brains.

## METHODS

### Participant samples

We evaluated complete GWAS results in the form of summary statistics (p-values and odds ratios) for clinically diagnosed AD dementia and seven CV associated RFs, including BMI [20], T2D [32], CAD [25], WHR [33], and plasma lipid levels (TC, LDL, and HDL [42]; Table 1). We obtained publicly available AD GWAS summary statistic data from the International Genomics of Alzheimer’s Disease Project (IGAP Stage 1, for additional details see Supplemental Information and [17]; Table 1). The IGAP Stage 1 consists of 17,008 AD cases (mean age = 74.7 ± 7.7 years; 59.4% female) and 37,154 controls (mean age = 76.3 ± 8.1 years; 58.6% female) drawn from four different consortia across North America and Europe with genotyped or imputed data at 7,055,881 SNPs (for a description of the AD dementia cases and controls within the IGAP Stage 1 sub-studies, please see [17]).

**Table 1.**
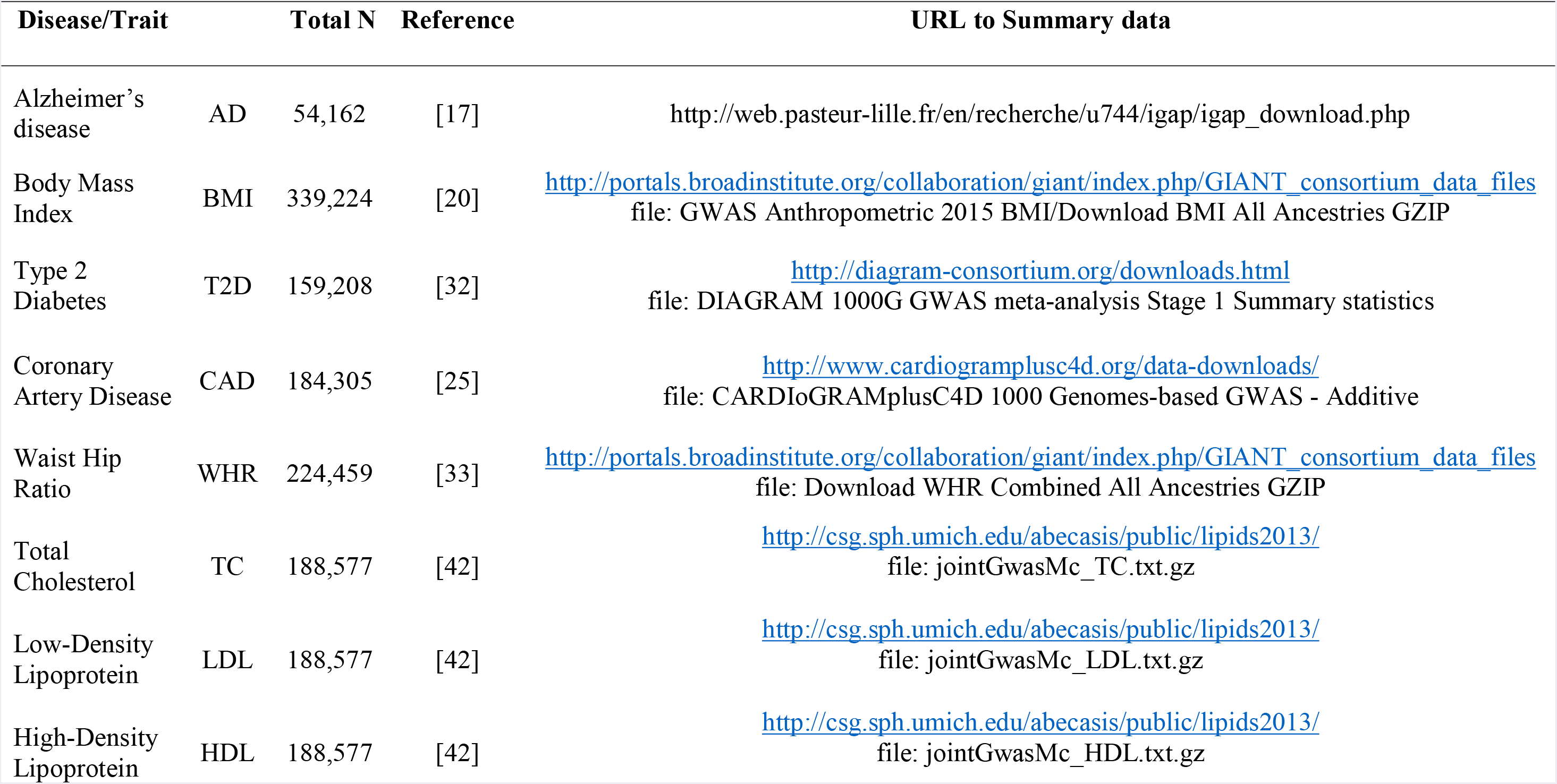
Summary data from all GWAS used in the current study

| Disease/Trait |  | Total N | Reference | URL to Summary data |
| --- | --- | --- | --- | --- |
| Alzheimer's disease | AD | 54,162 | [17] | http://web.pasteur-lille.fr/en/recherche/u744/igap/igap_download.php |
| Body Mass Index | BMI | 339,224 | [20] | [http://portals.broadinstitute.org/collaboration/giant/index.php/GIANT consortium data files](http://portals.broadinstitute.org/collaboration/giant/index.php/GIANT_consortium_data_files)<br>file: GWAS Anthropometric 2015 BMI/Download BMI All Ancestries GZIP |
| Type 2 Diabetes | T2D | 159,208 | [32] | http://diagram-consortium.org/downloads.html<br>file: DIAGRAM 1000G GWAS meta-analysis Stage 1 Summary statistics |
| Coronary Artery Disease | CAD | 184,305 | [25] | http://www.cardiogramplusc4d.org/data-downloads/<br>file: CARDIoGRAMplusC4D 1000 Genomes-based GWAS - Additive |
| Waist Hip Ratio | WHR | 224,459 | [33] | [http://portals.broadinstitute.org/collaboration/giant/index.php/GIANT consortium data files](http://portals.broadinstitute.org/collaboration/giant/index.php/GIANT_consortium_data_files)<br>file: Download WHR Combined All Ancestries GZIP |
| Total Cholesterol | TC | 188,577 | [42] | http://csg.sph.umich.edu/abecasis/public/lipids2013/<br>file: jointGwasMc_TC.txt.gz |
| Low-Density Lipoprotein | LDL | 188,577 | [42] | http://csg.sph.umich.edu/abecasis/public/lipids2013/<br>file: jointGwasMc_LDL.txt.gz |
| High-Density Lipoprotein | HDL | 188,577 | [42] | http://csg.sph.umich.edu/abecasis/public/lipids2013/<br>file: jointGwasMc_HDL.txt.gz |

We obtained publicly available BMI GWAS summary statistic data from the Genetic Investigation of Anthropometric Traits consortium (GIANT, for additional details see [20]; Table 1). The BMI GWAS meta-analysis consists of 339,224 individuals of European and non-European descent from 125 studies. The meta-analysis summary statistics were adjusted for age, sex and inverse normal transformation of the residuals.

The publicly available T2D GWAS summary statistic data were obtained from the Diabetes Genetics Replication and Meta-analysis consortium (DIAGRAM, for additional details see [32]; Table 1). The T2D GWAS meta-analysis Stage 1 summary statistics were drawn from 18 studies and consists of 26,676, T2D cases and 132,532 controls from European and non-European decent. Stage 1 summary statistics were adjusted for age, sex and principal components derived from the genetic data to account for population stratification. Genomic control (GC) correction to study-level was also conducted to correct for residual population structure not accounted for by principal components adjustment.

We obtained publicly available CAD GWAS summary statistic data from the CARDIoGRAMplusC4D consortium (Coronary Artery Disease Genome wide Replication and Meta-analysis (CARDIoGRAM) plus The Coronary Artery Disease (C4D) Genetics). The CARDIoGRAMplusC4D 1000 Genomes-based GWAS meta-analysis consists of 60,801 CAD cases and 123,504 controls from European and non-European decent (for additional details see [25]; Table 1). The meta-analysis statistics were adjusted for over-dispersion.

We obtained publicly available WHR without adjustment for BMI GWAS summary statistic data from the GIANT consortium (for additional details see [33]; Table 1). The WHR GWAS meta-analysis summary statistics included 224,459 individuals of European and non-European decent. The summary statistics were adjusted for age and study-specific covariates. Residuals were calculated for men and women separately and then transformed by the inverse standard normal function.

Lastly, we obtained publicly available TC, LDL, and HDL GWAS summary statistic data from Global Lipids Genetics Consortium (for additional details see [42]; Table 1). The TC, LDL, and HDL GWAS summary statistics included 188,577 individuals of European and non-European decent. The summary statistics were adjusted for age and sex.

The relevant institutional review boards or ethics committees approved the research protocol of all individual GWAS used in the current analysis, and all human participants gave written informed consent.

### Genetic Enrichment and Conjunction False Discovery Rates (FDR)

We evaluated whether there is pleiotropic enrichment in AD as a function of each of the seven CV RFs (see Supplemental Information). These validated methods have been described previously [2, 3, 6, 16, 44]. Briefly, for given associated phenotypes A (e.g. AD) and B (e.g. BMI), pleiotropic ‘enrichment’ of phenotype A with phenotype B exists if the proportion of SNPs or genes associated with phenotype A increases as a function of increased association with phenotype B. To assess for enrichment, we constructed fold-enrichment plots of nominal – log_10_(p) values for all AD SNPs and for subsets of SNPs determined by the significance of their association with each of the seven CV RFs. In fold-enrichment plots, the presence of enrichment is reflected as an upward deflection of the curve for phenotype A if the degree of deflection from the expected null line is dependent on the degree of association with phenotype B. To assess for polygenic effects below the standard GWAS significance threshold, we focused the fold-enrichment plots on SNPs with nominal –log_10_(p) < 7.3 (corresponding to p > 5×10^−8^). The enrichment seen can be directly interpreted in terms of true discovery rate (TDR = 1 – False Discovery Rate (FDR)) (for additional details see Supplemental Information). To account for large blocks of linkage disequilibrium (LD) that may result in spurious genetic enrichment, we applied a random pruning approach, where one random SNP per LD block (defined by an r^2^ of 0.8) was used and averaged over 200 random pruning runs. Given prior evidence that several genetic variants within the human leukocyte antigen (*HLA*) region on chromosome 6 [38, 44], microtubule-associated tau protein (*MAPT*) region on chromosome 17 [9] and the *APOE* region on chromosome 19 [10] are associated with increased AD risk, one concern is that random pruning may not sufficiently account for these large LD blocks resulting in artificially inflated genetic enrichment [6]. To better account for these large LD blocks, in our genetic enrichment analyses, we removed all SNPs in LD with r^2^ > 0.2 within 1Mb of *HLA, MAPT* and *APOE* variants (based on 1000 Genomes Project LD structure).

To identify specific loci jointly involved with AD and the seven CV RFs, we computed conjunction FDRs, as previously described [2, 3, 6, 43, 44]. Briefly, conjunction FDR, denoted by FDR _trait1 & trait2_ is defined as the posterior probability that a SNP is null for either trait or for both simultaneously, given the *p*-values for both traits are as small, or smaller, than the observed *p*-values. Unlike the *conditional* FDR which ranks disease/primary phenotype associated SNPs based on genetic ‘relatedness’ with secondary phenotypes, the *conjunction* FDR minimizes the possibility/likelihood of a single phenotype driving the common association signal. *Conjunction* FDR therefore is more conservative and specifically pinpoints pleiotropic loci between the traits of interest. We used an overall FDR threshold of < 0.05, which means 5 expected false discoveries per hundred reported. Manhattan plots were constructed based on the ranking of conjunction FDR to illustrate the genomic location of the pleiotropic loci. In all analyses, we controlled for the effects of genomic inflation by using intergenic SNPs (see Supplemental Information). Detailed information on fold enrichment plots, Manhattan plots, and conjunction FDR can be found in Supplemental Information and prior reports [2, 3, 6, 43, 44].

### Functional evaluation of shared risk loci

To assess whether SNPs that are shared between AD and CV RFs modify gene expression, we identified *cis*-expression quantitative loci (eQTLs, defined as variants within 1 Mb of a gene’s transcription start site) and regional brain expression of AD/CV SNPs in a publicly available dataset of normal control brains (UKBEC, http://braineac.org [30]). Given the evaluation of CV RFs, we also evaluated eQTLs using a blood-based dataset [41], We applied an analysis of covariance (ANCOVA) to test for associations between genotypes and gene expression. We tested SNPs using an additive model.

### Functional association analyses

To evaluate enrichment in tissue types and biological pathways of the AD/CV pleiotropic genes, we used FUMA, a web-based platform that integrates information from multiple biological resources to facilitate functional annotation of GWAS results [40]. To evaluate potential protein and genetic interactions, co-expression, co-localization and protein domain similarity between the overlapping genes, we used GeneMANIA, (http://genemania.org), an online web-portal for bioinformatic assessment of gene networks [39].

### Gene expression alterations in AD brains

To determine whether the AD/CV pleiotropic genes are differentially expressed in AD brains, we analyzed gene expression of overlapping genes in publicly available datasets. Mayo Clinic Brain Bank (Mayo) RNAseq study was accessed from the Accelerating Medicines Partnership – Alzheimer’s Disease (AMP-AD) portal (syn3163039; accessed April 2017). We examined gene expression in the temporal cortex of brains with neuropathologic diagnosis of AD dementia (N=82) and elderly control brains that lacked a diagnosis of neurodegenerative disease (n=80) [1]. Differential gene expression comparing AD dementia vs. controls used a “Simple Model.” In this model, multi-variable linear regression analyses were conducted in R, using CQN normalized gene expression measures and including age at death, gender, RNA integrity number (RIN), brain tissue source, and flowcell as biological and technical covariates.

### Evaluation of cell classes within the brain

Using a publicly available RNA-sequencing transcriptome and splicing database [46], we ascertained whether the AD/CV pleiotropic genes are expressed by specific cell classes within the brain. The cell types surveyed are neurons, fetal astrocytes, mature astrocytes, oligodendrocytes, microglia/macrophages, and endothelial cells (for additional details, see [46]).

## RESULTS

### Pleiotropic enrichment in AD conditional on plasma lipid levels

For progressively stringent p-value thresholds for AD SNPs (i.e. increasing values of nominal –log_10_(p)), we found at least 40-fold enrichment using TC, 20-fold enrichment using HDL and 60-fold enrichment using LDL (Figure 1). In comparison, we found no enrichment with BMI, T2D, CAD, and WHR. We note that these results reflect genetic enrichment in AD as a function of CV RFs after the exclusion of SNPs in LD with *HLA, MAPT*, and *APOE* (see Methods).

**Fig. 1.**
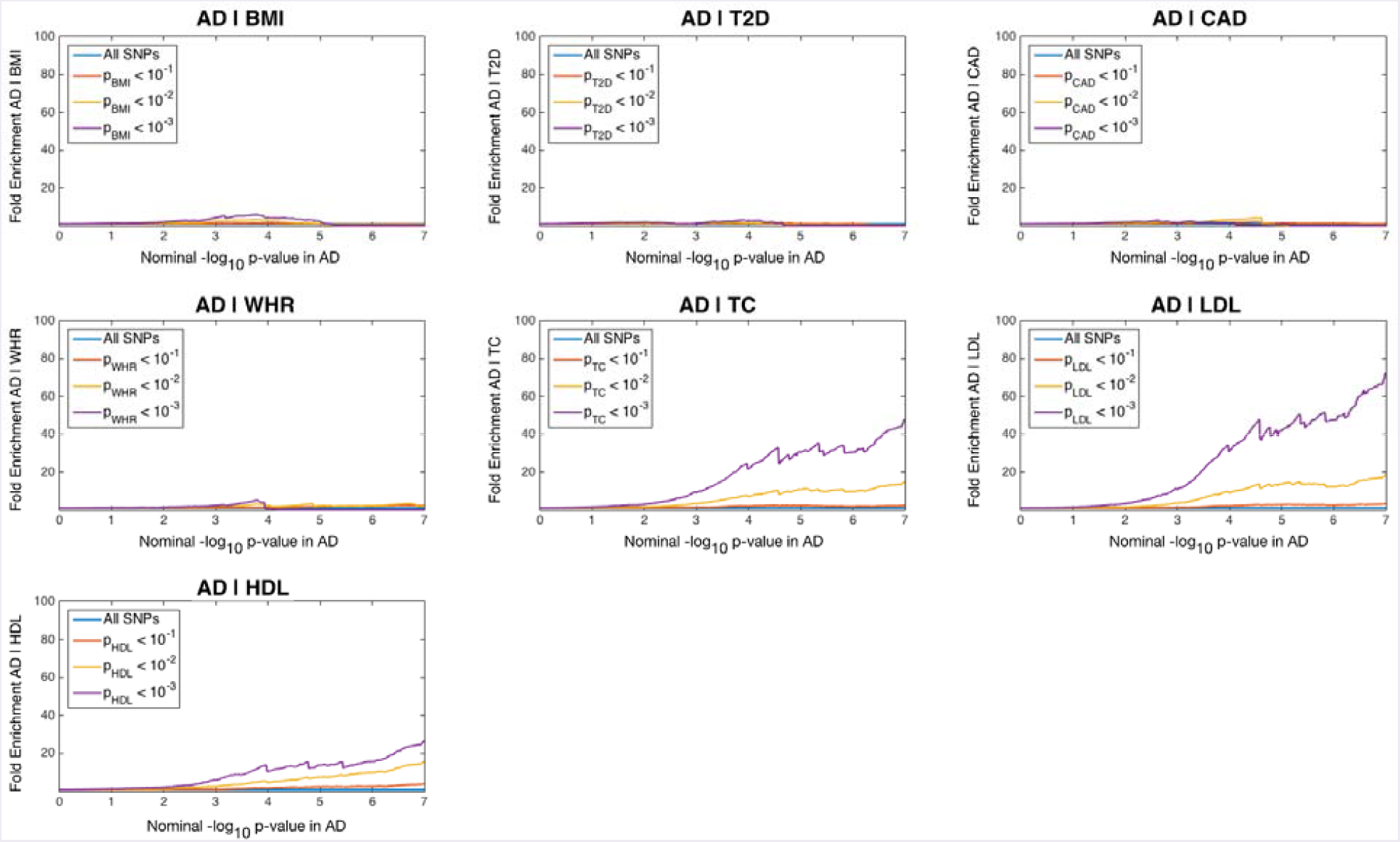
Fold enrichment plots of nominal -log_10_ p-values (corrected for inflation and excluding *APOE, MAPT*, and *HLA* regions) in Alzheimer’s disease (AD) below the standard GWAS threshold of *p* < 5×10^−8^ as a function of significance of association with body mass index (BMI), type 2 diabetes (T2D), coronary artery disease (CAD), waist hip ratio (WHR), total cholesterol (TC), low-density lipoprotein (LDL), and high-density lipoprotein (HDL) at the level of *p* ≤ 1, *p* ≤ 0.1, *p* ≤ 0.01, respectively. Blue line indicates all SNPs

Given the long-range LD associated with the *APOE/TOMM40* region [45], we focused our pleiotropy analyses on genetic variants outside chromosome 19. At a conjunction FDR < 0. 05, we identified 57 SNPs, in total, across 19 chromosomes jointly associated with AD and CV RFs (Figure 2, Table 2). Of these, we found that on chromosome 11, rs11039149 (closest gene = *NR1H3*) was in LD with: a) rs2071305 (closest gene = *MYBPC3*, pairwise D’ = 0.78) and b) rs4752856 (closest gene = *MTCH2*, pairwise D’ = 0.86). None of the remaining SNPs showed strong LD, suggesting that each SNP contributed independently to the genetic enrichment signal.

**Table 2.** Overlapping loci between AD and CV RFs at a conjunction FDR < 0.05. Chromosome 19 SNPs are excluded.

|  | SNP | Chr | Position | Nearest Gene | A1 | Reference<br>CV RF | Min<br>Conj FDR | Reference<br>CV RF<br>p-value | AD<br>p-value | Direction<br>of Allelic<br>Effect |
| --- | --- | --- | --- | --- | --- | --- | --- | --- | --- | --- |
| 1 | rs11580687 | 1 | 193685925 | CDC73 | G | BMI | 2.05E-02 | 2.46E-04 | 8.63E-05 | +/- |
| 2 | rs12410656 | 1 | 27397604 | SLC9A1 | T | LDL | 3.48E-03 | 9.32E-09 | 1.03E-04 | -/+ |
| 3 | rs6676563 | 1 | 55652983 | USP24 | G | LDL | 4.16E-02 | 2.12E-14 | 1.62E-03 | +/- |
| 4 | rs72796734 | 2 | 44063731 | ABCG5 | T | LDL | 2.38E-02 | 1.61E-04 | 8.41E-04 | -/+ |
| 5 | rs741477 | 2 | 65066311 | AK097952 | G | TC | 2.28E-02 | 7.96E-04 | 6.79E-04 | -/- |
| 6 | rs12994639 | 2 | 64959331 | SERTAD2 | G | TC | 4.80E-02 | 2.37E-06 | 1.61E-03 | -/- |
| 7 | rs17713879 | 2 | 254215 | SH3YL1 | A | HDL | 4.35E-02 | 8.03E-04 | 7.34E-04 | -/+ |
| 8 | rs6785930 | 3 | 151056616 | MED12L | A | LDL | 4.82E-02 | 3.17E-03 | 8.67E-04 | +/- |
| 9 | rs17037999 | 4 | 108613027 | PAPSS1 | G | LDL | 4.56E-02 | 2.98E-03 | 4.35E-04 | +/+ |
| 10 | rs6886253 | 5 | 174616451 | DRD1 | T | LDL | 1.85E-02 | 5.23E-04 | 6.36E-04 | -/- |
| 11 | rs5744712 | 5 | 74892002 | POLK | C | LDL | 3.61E-02 | 4.32E-32 | 1.36E-03 | +/+ |
| 12 | rs141974113 | 6 | 32360050 | BTNL2 | A | CAD | 3.20E-02 | 2.64E-04 | 2.06E-04 | +/+ |
| 13 | rs646984 | 6 | 32576592 | HLA-DRB5 | C | TC | 2.14E-03 | 1.27E-04 | 2.06E-05 | -/+ |
| 14 | rs2717926 | 7 | 37714304 | BC043356 | C | BMI | 1.93E-02 | 2.26E-04 | 9.55E-05 | -/- |
| 15 | rs2888877 | 7 | 92228400 | CDK6 | T | LDL | 3.18E-02 | 1.31E-03 | 1.00E-03 | -/- |
| 16 | rs17153037 | 7 | 26048075 | DM004234 | A | BMI | 3.21E-02 | 1.02E-04 | 1.85E-04 | +/- |
| 17 | rs35991721 | 7 | 99728790 | MBLAC1 | T | CAD | 1.14E-02 | 5.28E-05 | 5.92E-05 | -/+ |
| 18 | rs702483 | 7 | 6426941 | RAC1 | T | TC | 2.95E-02 | 1.90E-03 | 6.27E-04 | -/+ |
| 19 | rs858513 | 7 | 99825558 | STAG3OS | G | LDL | 1.63E-02 | 4.95E-04 | 5.51E-04 | -/+ |
| 20 | rs7819800 | 8 | 95975168 | C8ORF38 | A | T2D | 3.81E-02 | 1.68E-04 | 1.24E-04 | -/+ |
| 21 | rs10093964 | 8 | 27312040 | PTK2B | T | BMI | 3.15E-02 | 4.29E-04 | 1.13E-04 | +/- |
| 22 | rs7014168 | 8 | 10641965 | SOX7 | A | HDL | 2.80E-02 | 9.71E-11 | 4.35E-04 | -/- |
| 23 | rs1180628 | 8 | 116430861 | TRPS1 | C | LDL | 3.67E-02 | 1.24E-04 | 1.39E-03 | +/- |
| 24 | rs1883025 | 9 | 107664301 | ABCA1 | T | LDL | 4.28E-03 | 9.42E-12 | 1.29E-04 | +/- |
| 25 | rs943424 | 9 | 137436967 | COL5A1 | T | TC | 3.10E-02 | 2.17E-03 | 1.87E-04 | +/+ |
| 26 | rs7916835 | 10 | 124122373 | PLEKHA1 | T | T2D | 1.44E-02 | 1.62E-05 | 4.41E-05 | -/- |
| 27 | rs1263173 | 11 | 116681008 | APOA4 | G | TC | 4.59E-02 | 5.84E-07 | 1.53E-03 | +/+ |
| 28 | rs2727273 | 11 | 61702603 | BEST1 | T | LDL | 2.37E-02 | 6.56E-04 | 8.40E-04 | -/- |
| 29 | rs7928842 | 11 | 47566352 | CELF1 | C | BMI | 1.82E-02 | 8.72E-11 | 8.85E-05 | -/+ |
| 30 | rs4752856 | 11 | 47648042 | MTCH2 | A | TC | 4.36E-04 | 6.79E-07 | 9.14E-06 | -/- |
| 31 | rs2071305 | 11 | 47370957 | MYBPC3 | C | HDL | 3.10E-04 | 1.29E-10 | 3.09E-06 | -/- |
| 32 | rs11039149 | 11 | 47276675 | NR1H3 | G | LDL | 2.21E-02 | 1.38E-03 | 4.97E-06 | +/+ |
| 33 | rs12292911 | 11 | 47449072 | PSMC3 | A | BMI | 1.19E-03 | 1.72E-07 | 3.78E-06 | +/- |
| 34 | rs3815364 | 11 | 67202847 | RPS6KB2 | A | LDL | 3.82E-02 | 6.36E-04 | 1.46E-03 | +/- |
| 35 | rs11038670 | 11 | 45845150 | SLC35C1 | A | HDL | 2.29E-02 | 2.02E-04 | 3.43E-04 | +/+ |
| 36 | rs2510044 | 11 | 77909014 | USP35 | A | TC | 4.68E-02 | 6.09E-05 | 1.57E-03 | -/- |
| 37 | rs12146727 | 12 | 7170336 | C1S | A | CAD | 2.20E-02 | 1.63E-04 | 9.30E-05 | -/- |
| 38 | rs10861632 | 12 | 106960182 | RFX4 | A | HDL | 2.93E-02 | 2.45E-05 | 4.58E-04 | +/- |
| 39 | rs7972529 | 12 | 112841242 | RPL6 | G | CAD | 4.93E-02 | 4.05E-04 | 3.58E-04 | +/- |
| 40 | rs10773111 | 12 | 125332955 | SCARB1 | T | HDL | 4.94E-02 | 2.40E-12 | 8.55E-04 | +/+ |
| 41 | rs17101733 | 14 | 75123026 | AREL1 | G | TC | 6.54E-03 | 4.17E-04 | 3.52E-06 | +/+ |
| 42 | rs6493386 | 15 | 50213929 | ATP8B4 | G | LDL | 3.76E-02 | 1.46E-03 | 1.13E-03 | -/- |
| 43 | rs11857703 | 15 | 63379592 | TPM1 | A | HDL | 4.16E-02 | 7.63E-07 | 6.96E-04 | +/+ |
| 44 | rs3098197 | 15 | 50817133 | USP50 | A | HDL | 1.43E-02 | 1.95E-05 | 2.00E-04 | -/- |
| 45 | rs889555 | 16 | 31122571 | BCKDK | T | TC | 3.16E-02 | 1.04E-03 | 9.63E-04 | -/+ |
| 46 | rs8062895 | 16 | 72048632 | DHODH | G | LDL | 4.07E-02 | 2.79E-05 | 1.58E-03 | +/- |
| 47 | rs9941245 | 16 | 19916895 | GPRC5B | G | HDL | 1.63E-02 | 1.09E-04 | 2.32E-04 | -/+ |
| 48 | rs113260531 | 17 | 5138980 | BC029580 | A | CAD | 3.19E-02 | 2.66E-04 | 6.84E-06 | +/- |
| 49 | rs3760372 | 17 | 45380002 | ITGB3 | T | LDL | 1.31E-02 | 5.96E-07 | 4.32E-04 | +/- |
| 50 | rs144939739 | 17 | 43666820 | LOC644172 | A | CAD | 3.26E-02 | 2.01E-05 | 2.10E-04 | -/+ |
| 51 | rs12940610 | 17 | 64312463 | PRKCA | A | LDL | 2.25E-02 | 9.00E-04 | 7.93E-04 | -/+ |
| 52 | rs12951873 | 17 | 47403553 | ZNF652 | C | CAD | 3.04E-02 | 1.67E-04 | 1.93E-04 | +/+ |
| 53 | rs11083394 | 18 | 27940854 | DSC2 | G | LDL | 1.92E-02 | 2.81E-04 | 6.62E-04 | -/- |
| 54 | rs17656498 | 18 | 47128176 | LIPG | C | TC | 1.39E-02 | 3.02E-09 | 3.90E-04 | -/+ |
| 55 | rs6021721 | 20 | 50746651 | ZFP64 | G | HDL | 4.51E-02 | 9.01E-05 | 7.68E-04 | +/- |
| 56 | rs9621715 | 22 | 21942007 | UBE2L3 | A | BMI | 3.23E-02 | 3.66E-04 | 1.86E-04 | -/+ |
| 57 | rs2298428 | 22 | 21982892 | YDJC | T | TC | 3.70E-03 | 6.15E-06 | 9.17E-05 | -/- |

**Fig. 2.**
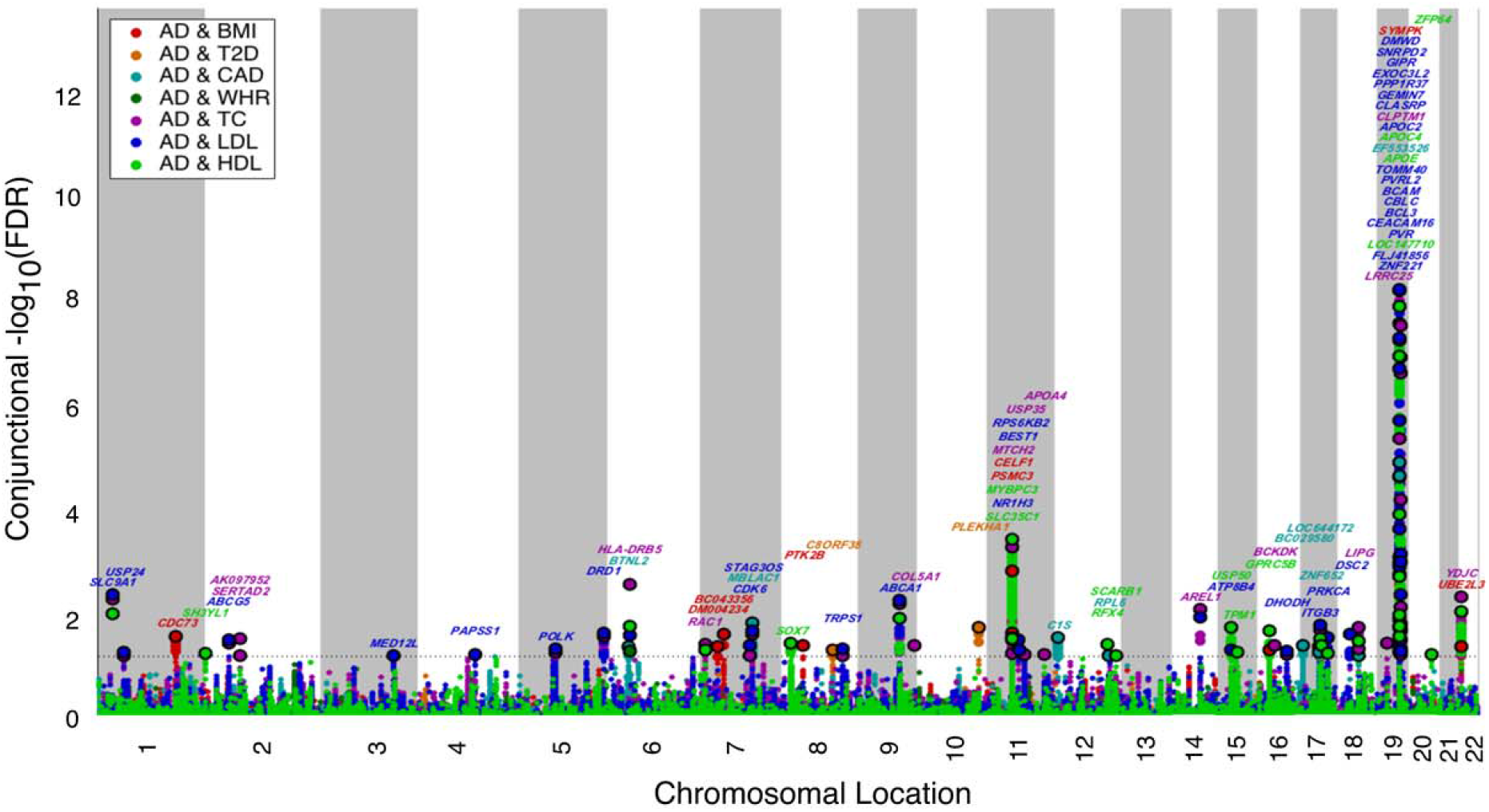
Conjunction Manhattan plot of conjunction –log_10_ (FDR) values for Alzheimer’s disease (AD) alone (black) and AD given body mass index (BMI; AD&BMI, red), type 2 diabetes (T2D; AD&T2D, orange), coronary artery disease (CAD; AD&CAD, aquamarine), waist hip ratio (WHR; AD&WHR, green), total cholesterol (TC; AD&TC, purple), low-density lipoprotein (LDL; &LDL, blue) and high-density lipoprotein (HDL, AD|HDL, bright green). SNPs with conjunction –log_10_ FDR > 1.3 (i.e. FDR < 0.05) are shown with large points. A black line around the large points indicates the most significant SNP in each LD block and this SNP was annotated with the closest gene, which is listed above

Outside of the known association with the *APOE/TOMM40* region on chromosome 19, we identified several AD/CV associated loci including: 1) rs11580687 (chr 1, nearest gene = *CDC73*, conditioning trait = BMI), 2) rs72796734 (chr 2, nearest gene = *ABCG5*, conditioning trait = LDL), 3) rs1883025 (chr 9, nearest gene = *ABCA1*, conditioning trait = LDL), 4) rs1263173 (chr 11, nearest gene = *APOA4*, conditioning trait = TC), and 5) rs6493386 (chr 15, nearest gene = *ATP8B4*, conditioning trait = LDL) (Table 2). On chromosome 11, we found several SNPs tagging genes within the *MTCH2/SPI1* locus that may be separate from *CELF1/CUGBP1* and *APOA4* (Supplemental Figure 1a-j and eQTL section below).

### Shared genetic risk between CV RFs

To evaluate whether the AD susceptibility loci listed in Table 2 are associated with a single CV RF or with multiple associated RFs, we constructed a matrix plot. For each of the 7 CV RFs, we plotted the magnitude and direction of associated z-scores for all 57 AD/CV SNPs/closest genes (Figure 3, Supplemental Table 1). We found that many of the AD/CV SNPs/closest genes were associated with multiple CV RFs and with different direction of effects. For example, a) rs1883025/ABCA1 was associated with a z-score of -6.82 with LDL (p-value of 9.42 x10^−12^), and -18.1 with HDL (p-value of 5.16 x 10^−73^) and b) rs4752856/MTCH2 was associated with a z-score of -10.03 with HDL (p-value of 1.02 x10^−23^), -4.97 with TC (p-value of 6.79 x10^−7^) and +7.92 with BMI (p-value of 2.41 x 10^−15^). These findings illustrate that common genetic variants influencing AD risk are associated with multiple CV RFs.

**Fig. 3.**
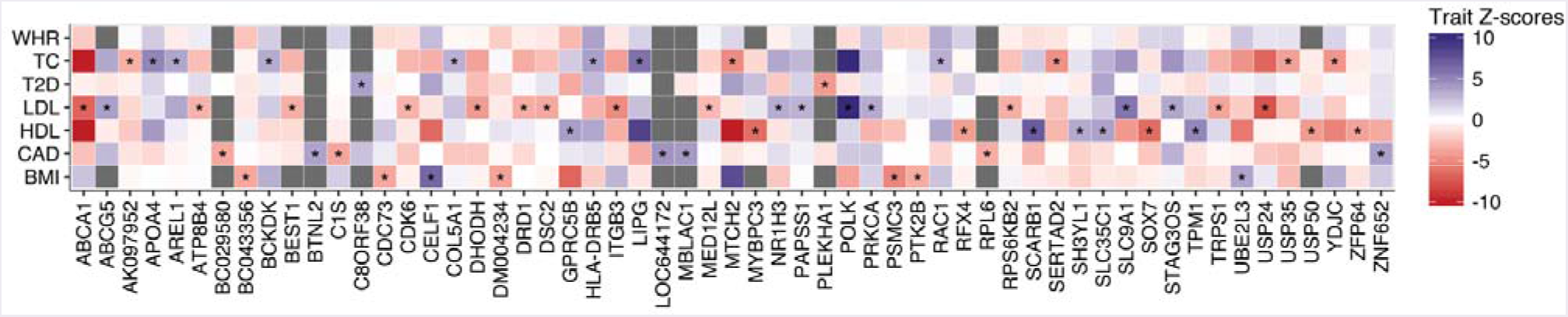
Matrix plot mapping directionally encoded z-scores for the *non-APOE* AD/CV pleiotropic genes for each CV RF. * indicates the conditioning RF used to identify the most significant SNP (see Table 2 and Fig. 2)

### cis-eQTLs

We found significant *cis*-associations between the AD/CV pleiotropic SNPs and 173 genes in either brain or blood tissue types (Supplemental Table 2). Several SNPs showed significant *cis*-eQTLs with multiple genes. Outside of the *APOE/TOMM40* region, we found 12 *cis*-eQTLs that replicated in both datasets, namely *C1QTNF4, CRY2, DMWD, HLA-DOB, KLC3, LACTB, MTCH2, NARS2, NUP160, PAPSS1, PTK2B*, and *SPI1*. Within the *APOE/TOMM40* region, we found a *cis*-association between rs3852860 and *PVRL2* that was significant in both brain and blood suggesting AD/CV pleiotropic genetic signal on chromosome 19 that is independent from *APOE.*

### Gene ontology and gene-protein network analyses

At an FDR < 0.05, we found that the closest genes associated with the AD/CV polymorphisms were associated with multiple gene ontology (GO) molecular, cellular, and biological pathways and tissue types. These shared loci were highly enriched for lipid-associated processes including cholesterol and sterol transport, regulation, binding, and activity (Supplemental Figure 2). Beyond *APOE*, several genes were associated with lipid metabolism including *ABCA1, ATP8B4, APOA4, ABCG5, ITGB3, LIPG*, and *NR1H3.* We also found that expression of the AD/CV pleiotropic genes was over-represented within different tissue types, in particular the cerebral cortex, skin, subcutaneous fat and arterial structures including the coronary artery (Supplemental Figure 3a). Beyond *APOE*, several genes were highly expressed across a number of tissue types including *C1S, PSMC3, PTK2B, RAC1*, and *RPL6* (Supplemental Figure 3b). In our ‘network’ analysis, we found that the majority of the AD/CV pleiotropic genes were co-expressed (53%), co-localized (13%), or showed shared protein domains (10%) (Figure 4). Functional analyses across functionally expressed AD/CV pleiotropic genes (i.e. those with significant *cis*-eQTLs in both brain and blood) are shown in Supplemental Figures 4-6.

**Fig. 4.**
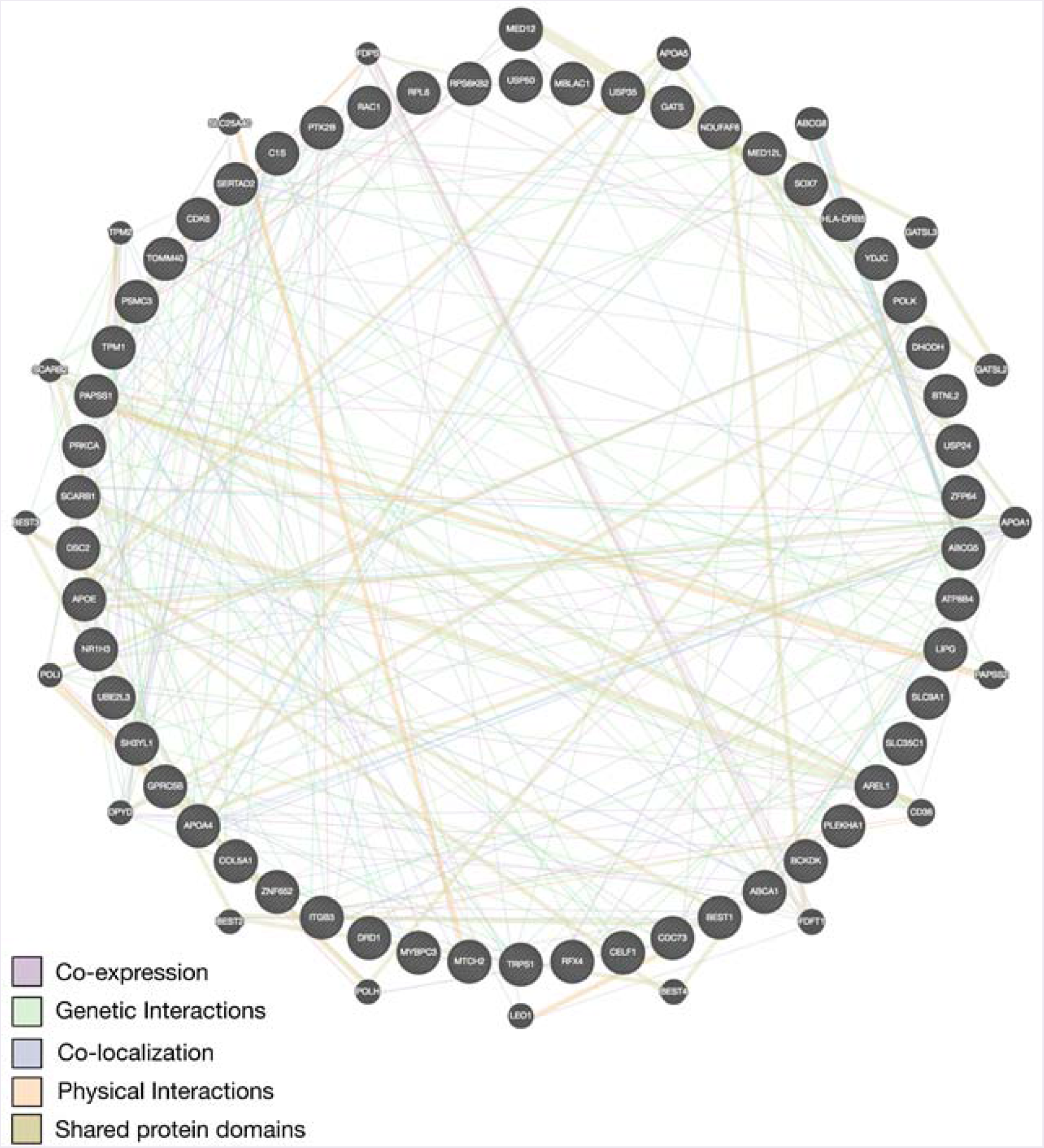
Network interaction graph predominantly illustrating co-expressed (purple), co-localized (blue), and shared protein domains (khaki) for AD/CV pleiotropic genes

### Gene expression in brains from AD patients and healthy controls

To investigate whether the AD/CV pleiotropic genes are differentially expressed in AD brains, we compared gene expression in AD brains with neuropathologically normal control brains. For several shared AD/CV genes, we observed differential expression in AD brains compared to controls (Supplemental Table 3). We found the strongest effects (absolute magnitude of beta-coefficients) for differential expression of *CDC73, TRPS1, ABCA1, USP24*, and *ABCG5* (Figure 5). Similarly, across functionally expressed pleiotropic genes (i.e. those with significant *cis*-eQTLs in both brain and blood), we observed differential expression in AD brains (Figure 5, Supplemental Table 4).

**Fig. 5.**
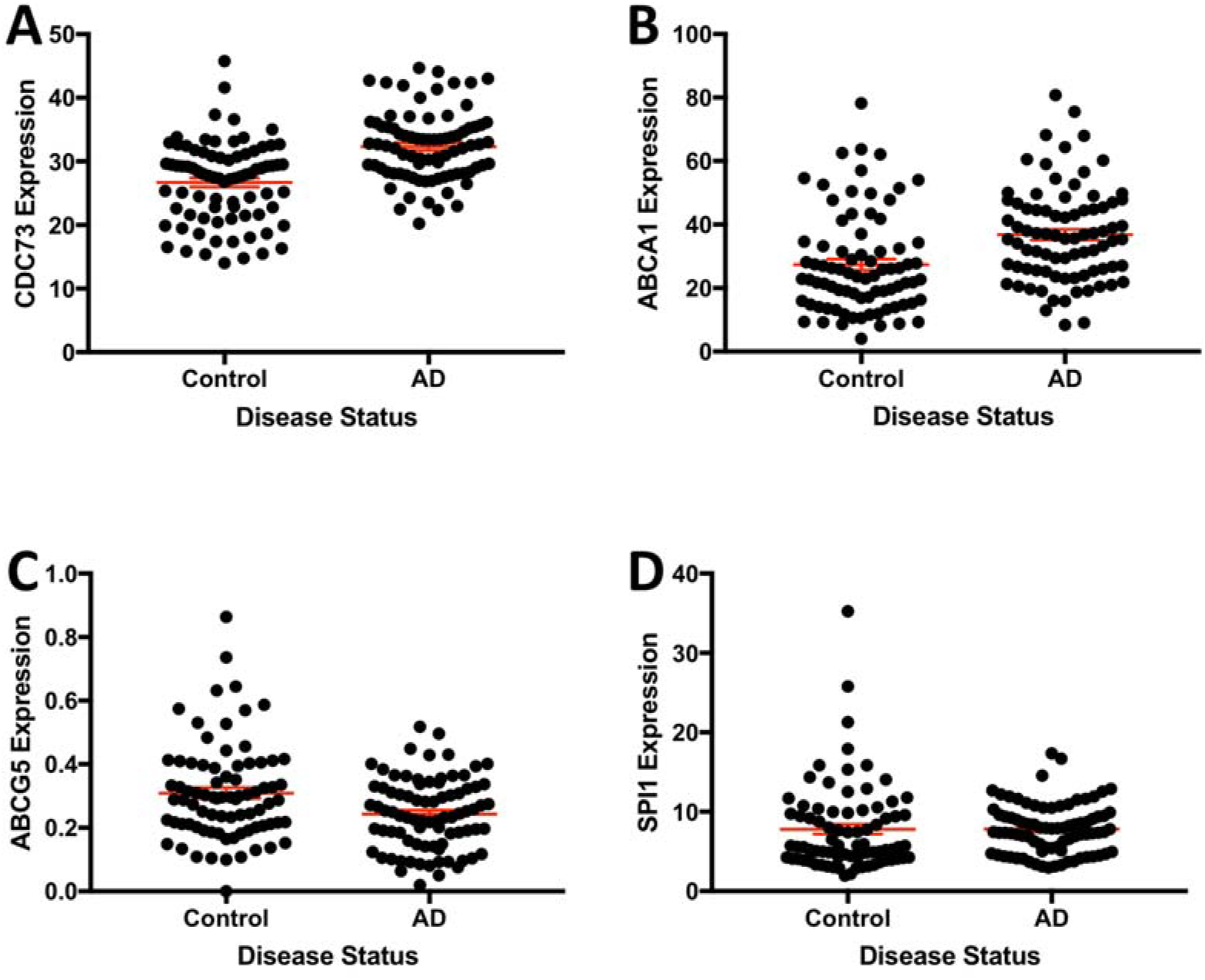
Genes associated with Alzheimer’s disease and cardiovascular disease differentially expressed in AD patients versus controls: a) CDC73, b) ABCA1, c) ABCG5, and d) SPI1

### Cell type enrichment

Across different central nervous system cell types, we found that the AD/CV pleiotropic genes were highly expressed in astrocytes (Figure 6). Similarly, across functionally expressed pleiotropic genes (i.e. those with significant *cis*-associations in both brain and blood), we found enrichment in astrocytes and microglia/macrophages (Figure 6). Notable examples include *ABCA1* (astrocytes), *MTCH2* (astrocytes), and *SPI1* (microglia/macrophages) (Supplemental Figures 7-8).

**Fig. 6.**
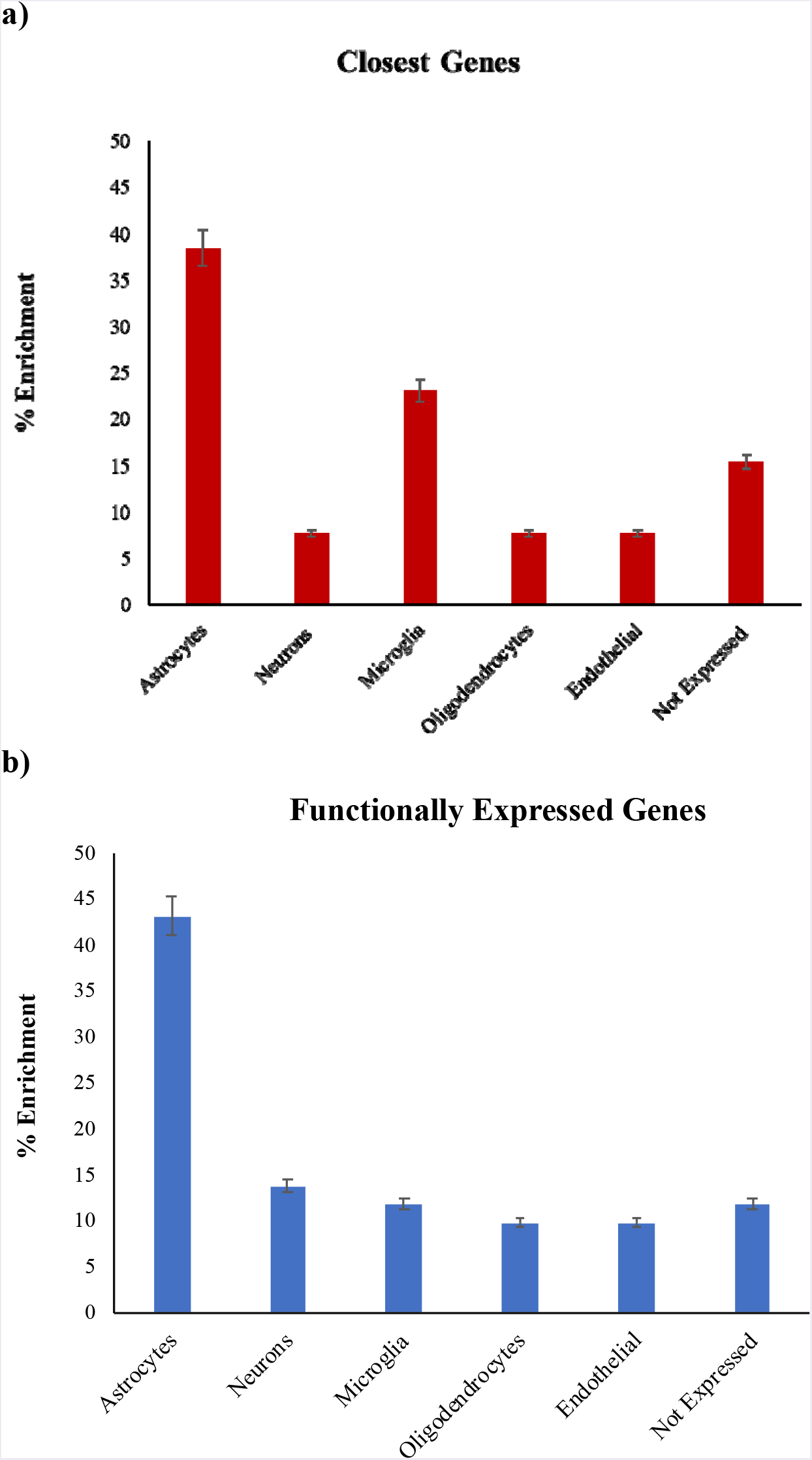
Enrichment associated with a) AD/CV pleiotropic genes in astrocytes and b) functionally expressed pleiotropic genes in astrocytes and microglia/macrophages. Functionally expressed pleiotropic genes were defined as genes with significant *cis*-associations in both brain and blood?

## DISCUSSION

Beyond *APOE*, we identified 57 SNPs on 19 different chromosomes that jointly conferred increased risk for AD and cardiovascular outcomes. Expression of these AD/CV pleiotropic genes was enriched for lipid and cholesterol metabolism processes, over-represented within the CNS (mainly astrocytes) and vascular structures, highly co-expressed, and differentially altered within AD brains. Collectively, our findings suggest that the polygenic component of AD is highly enriched for lipid associated RFs.

In their genetic association with AD, not all cardiovascular RFs are created equal. We found minimal genetic enrichment in AD as a function of T2D, BMI, WHR, and CAD suggesting that the known comorbidity [21, 28, 35] between these CV RFs and Alzheimer’s etiology is likely not genetic. In contrast, genetic enrichment in AD was predominantly localized to plasma lipids. Building on our prior work leveraging statistical power from large CV GWASs for AD gene discovery [10], we found genetic variants jointly associated with AD and CV RFs, many with known cholesterol/lipid function. By conditioning on plasma TC, LDL, and HDL levels, we identified AD susceptibility loci within genes encoding apolipoproteins, such as *APOA4*, ATP-binding cassette transporters, such as *ABCA1* and *ABCG5*, and phospholipases, such as *ATP8B4* and *LIPG* (for a discussion on lipid genes and AD see [11]). In functional analyses, expression of these pleiotropic genes was enriched for lipid metabolism pathways, over-represented within the brain and arteries, and perturbed within the brains of AD patients. Considered together with our co-expression findings, these results are consistent with the hypothesis that a network of non-*APOE* genes implicated in lipid biology also influence Alzheimer’s pathobiology.

Our pleiotropy findings suggest that complex diseases and traits have a complex genetic architecture. Although we did not evaluate causal associations using a Mendelian Randomization (MR) framework, our results have implications for the relationship between common genetic variants, cardiovascular RFs and AD as an outcome. To date, MR studies have typically evaluated a single CV risk factor at a time, which is valid only if the genetic variants used for the MR influence AD exclusively via the selected CV risk factor [18, 27]. For a number of variants, we found pleiotropy challenging the conventional MR approach (for a review on the assumptions underlying MR see [12]); that is, common genetic variants influencing AD are associated with multiple CV RFs, at times with a different directionality of effect (Figure 3, Supplemental Table 2). Instead of a single causal link between genetic variants, RF and the outcome [13], these results suggest two possible scenarios: 1. genetic variants influence cardiovascular RFs and AD independently, or, 2. genetic variants influence AD through multiple cardiovascular RFs.

On chromosome 11, our pleiotropy and eQTL results point to AD associated genetic signal within *MTCH2* and *SPI1*, independent from *CELF1/CUGB1.* By conditioning on cardiovascular RFs, we identified several SNPs tagging variants within *NR1H3, MTCH2, MYBPC3* and *CELF1.* The eQTL analyses showed *cis*-associations with *MTCH2* and *SPI1*, both of which showed differential gene expression alterations in AD brains. Consistent with our findings, a recent study found an AD risk locus within the *CELF1* region that was associated with lower expression of *SPI1* in monocytes and macrophages [14, 15]. *SPI1* encodes a transcription factor, PU.1, that is essential for myeloid cell development and a major regulator of cellular communication in the immune system [23]. We note that the majority of our pleiotropic genes were expressed in astrocytes and to a lesser extent, microglia, implicating genes expressed in microglia, astrocytes or other myeloid cell types in AD pathogenesis [34].

Our findings have clinical implications. First, given the common co-occurrence of vascular and Alzheimer’s pathology, it is highly likely that the clinically diagnosed AD individuals from our cohort have concomitant vascular brain disease, which may further contribute to their cognitive decline and dementia. As such, a plausible interpretation of our findings is that the susceptibility loci identified in this study may increase brain vulnerability to vascular and/or inflammatory insults, which in turn may exacerbate the clinical consequences of AD pathological changes. Second, no single common variant detected in this study will be clinically informative. Rather, integration of these pleiotropic variants into a cardiovascular pathway specific, polygenic ‘hazard’ framework for predicting AD age of onset may help identify older individuals jointly at risk for cardiovascular and Alzheimer’s disease [8]. Therapeutically targeting cardiovascular RFs in these individuals may impact the Alzheimer’s disease trajectory.

This study has limitations. First, whereas several of the CV RF GWASs included European and non-European ancestry individuals, the IGAP AD cohort restricted analyses to non-Hispanic Whites. Therefore, these results may not be generalizable to AD patients from other populations. Second, our AD patients were diagnosed largely using clinical criteria without neuropathology confirmation and this may result in misclassification of case status. However, such misclassification should reduce statistical power and bias results toward the null. Finally, given evidence that phospholipids are proinflammatory [29], future work should evaluate whether LDL, HDL or TC influence AD risk through inflammation or other mediator variables.

In summary, across a large cohort (n > 500,000 cases and controls), we show cardiovascular associated polygenic enrichment in AD. Beyond *APOE*, our findings support a disease model in which lipid biology is integral to the development of clinical AD in a subset of individuals. Lastly, considerable clinical, pathological and epidemiological evidence has shown overlap between Alzheimer’s and cardiovascular risk factors; here, we provide genetic support for this association.

## ACKNOWLEDGEMENTS

We thank the Shiley-Marcos Alzheimer’s Disease Research Center at UCSD and the Memory and Aging Center at UCSF for continued support and the International Genomics of Alzheimer’s Project (IGAP) for providing summary results data for these analyses. This work was supported by grants from the National Institutes of Health (NIH-AG046374, K01AG049152), National Alzheimer’s Coordinating Center Junior Investigator Award (RSD), RSNA Resident/Fellow Award (RSD), Foundation ASNR Alzheimer’s Imaging Grant (RSD), the Research Council of Norway (#213837, #225989, #223273, #237250/EU JPND), the South East Norway Health Authority (2013-123), Norwegian Health Association and the KG Jebsen Foundation.

## Declaration of interests

JBB served on advisory boards for Elan, Bristol-Myers Squibb, Avanir, Novartis, Genentech, and Eli Lilly and holds stock options in CorTechs Labs, Inc. and Human Longevity, Inc. AMD is a founder of and holds equity in CorTechs Labs, Inc., and serves on its Scientific Advisory Board. He is also a member of the Scientific Advisory Board of Human Longevity, Inc. (HLI), and receives research funding from General Electric Healthcare (GEHC). The terms of these arrangements have been reviewed and approved by the University of California, San Diego in accordance with its conflict of interest policies.

## REFERENCES

1. Allen M, Carrasquillo MM, Funk C, et al. Human whole genome genotype and transcriptome data for Alzheimer’s and other neurodegenerative diseases. Scientific Data. 2016;3:160089. doi:10.1038/sdata.2016.89

2. Andreassen OA, Thompson WK, Dale AM. Boosting the Power of Schizophrenia Genetics by Leveraging New Statistical Tools. Schizophr Bull. 2014;40(1):13–17. doi:10.1093/schbul/sbt168

3. Andreassen OA, Thompson WK, Schork AJ, et al. Improved Detection of Common Variants Associated with Schizophrenia and Bipolar Disorder Using Pleiotropy-Informed Conditional False Discovery Rate. PLOS Genetics. 2013;9(4):e1003455. doi:10.1371/journal.pgen.1003455

4. Attems J, Jellinger KA. The overlap between vascular disease and Alzheimer’s disease--lessons from pathology. BMC Med. 2014;12:206. doi:10.1186/s12916-014-0206-2

5. Barnes DE, Yaffe K. The Projected Impact of Risk Factor Reduction on Alzheimer’s Disease Prevalence. Lancet Neurol. 2011;10(9):819–828. doi:10.1016/S1474-4422(11)70072-2

6. Broce I, Karch CM, Wen N, et al. Immune-related genetic enrichment in frontotemporal dementia: An analysis of genome-wide association studies. PLoS Med. 2018;15(1):e1002487. doi:10.1371/journal.pmed.1002487

7. Carmona S, Hardy J, Guerreiro R. Chapter 26 - The genetic landscape of Alzheimer disease. In: Geschwind DH, Paulson HL, Klein C, eds. Handbook of Clinical Neurology. Vol 148. Neurogenetics, Part II. Elsevier; 2018:395–408. doi:10.1016/B978-0-444-64076-5.00026-0

8. Desikan RS, Fan CC, Wang Y, et al. Genetic assessment of age-associated Alzheimer disease risk: Development and validation of a polygenic hazard score. PLOS Medicine. 2017;14(3):e1002258. doi:10.1371/journal.pmed.1002258

9. Desikan RS, Schork AJ, Wang Y, et al. Genetic overlap between Alzheimer’s disease and Parkinson’s disease at the MAPT locus. Mol Psychiatry. 2015;20(12):1588–1595. doi:10.1038/mp.2015.6

10. Desikan RS, Schork AJ, Wang Y, et al. Polygenic Overlap Between C-Reactive Protein, Plasma Lipids, and Alzheimer Disease. Circulation. 2015;131(23):2061–2069. doi:10.1161/CIRCULATIONAHA.115.015489

11. DiPaolo G, Kim T-W. Linking lipids to Alzheimer’s disease: cholesterol and beyond. Nat Rev Neurosci. 2011;12(5):284–296. doi:10.1038/nrn3012

12. Emdin CA, Khera AV, Kathiresan S. Mendelian Randomization. JAMA. 2017;318(19):1925–1926. doi:10.1001/jama.2017.17219

13. Emdin CA, Khera AV, Natarajan P, et al. Genetic Association of Waist-to-Hip Ratio With Cardiometabolic Traits, Type 2 Diabetes, and Coronary Heart Disease. JAMA. 2017;317(6):626–634. doi:10.1001/jama.2016.21042

14. Huang K-L, Marcora E, Pimenova AA, et al. A common haplotype lowers PU. 1 expression in myeloid cells and delays onset of Alzheimer’s disease. Nat Neurosci. 2017;20(8):1052–1061. doi:10.1038/nn.4587

15. Karch CM, Ezerskiy LA, Bertelsen S, Consortium (ADGC) ADG, Goate AM. Alzheimer’s Disease Risk Polymorphisms Regulate Gene Expression in the ZCWPW1 and the CELF1 Loci. PLOS ONE. 2016;11(2):e0148717. doi:10.1371/journal.pone.0148717

16. Karch CM, Wen N, Fan CC, et al. Selective Genetic Overlap Between Amyotrophic Lateral Sclerosis and Diseases of the Frontotemporal Dementia Spectrum. JAMA Neurol. 2018;75(7):860–875. doi:10.1001/jamaneurol.2018.0372

17. Lambert JC, Ibrahim-Verbaas CA, Harold D, et al. Meta-analysis of 74,046 individuals identifies 11 new susceptibility loci for Alzheimer’s disease. Nat Genet. 2013;45(12):1452–1458. doi:10.1038/ng.2802

18. Larsson SC, Traylor M, Malik R, et al. Modifiable pathways in Alzheimer’s disease: Mendelian randomisation analysis. BMJ. 2017;359:j5375.

19. Livingston G, Sommerlad A, Orgeta V, et al. Dementia prevention, intervention, and care. Lancet. 2017;390(10113):2673–2734. doi:10.1016/S0140-6736(17)31363-6

20. Locke AE, Kahali B, Berndt SI, et al. Genetic studies of body mass index yield new insights for obesity biology. Nature.2015;518(7538):197–206. doi:10.1038/nature14177

21. Luchsinger JA, Tang MX, Stern Y, Shea S, Mayeux R. Diabetes mellitus and risk of Alzheimer’s disease and dementia with stroke in a multiethnic cohort. Am J Epidemiol. 2001;154(7):635–641.

22. Mahley RW. Central Nervous System Lipoproteins: ApoE and Regulation of Cholesterol Metabolism. Arterioscler Thromb Vasc Biol. 2016;36(7):1305–1315. doi:10.1161 /ATVBAHA.116.307023

23. McKercher SR, Torbett BE, Anderson KL, et al. Targeted disruption of the PU.1 gene results in multiple hematopoietic abnormalities. EMBO J. 1996;15(20):5647–5658.

24. National Academies of Sciences, Engineering, and Medicine, Health and Medicine Division, Board on Health Sciences Policy, Committee on Preventing Dementia and Cognitive Impairment. Preventing Cognitive Decline and Dementia: A Way Forward. (Downey A, Stroud C, Landis S, Leshner AI, eds.). Washington (DC): National Academies Press (US); 2017. http://www.ncbi.nlm.nih.gov/books/NBK436397/. Accessed April 17, 2018.

25. Nikpay M, Goel A, Won H-H, et al.A comprehensive 1000 Genomes–based genome-wide association meta-analysis of coronary artery disease. Nature Genetics. 2015;47(10):1121–1130. doi:10.1038/ng.3396

26. Norton S, Matthews FE, Barnes DE, Yaffe K, Brayne C.Potential for primary prevention of Alzheimer’s disease: an analysis of population-based data. Lancet Neurol. 2014;13(8):788–794. doi:10.1016/S1474-4422(14)70136-X

27. Østergaard SD, Mukherjee S, Sharp SJ, et al.Associations between Potentially Modifiable Risk Factors and Alzheimer Disease: A Mendelian Randomization Study. PLoS Med. 2015;12(6):e1001841; doi:10.1371/journal.pmed.1001841

28. Profenno LA, Porsteinsson AP, Faraone SV. Meta-analysis of Alzheimer’s disease risk with obesity, diabetes, and related disorders. Biol Psychiatry. 2010;67(6):505–512. doi:10.1016/j.biopsych.2009.02.013

29. Que X, Hung M-Y, Yeang C, et al.Oxidized phospholipids are proinflammatory and proatherogenic in hypercholesterolaemic mice. Nature. 2018;558(7709):301–306. doi:10.1038/s41586-018-0198-8.

30. Ramasamy A, Trabzuni D, Guelfi S, et al. Genetic variability in the regulation of gene expression in ten regions of the human brain. Nature Neuroscience. 2014;17(10):1418–1428. doi:10.1038/nn.3801

31. Reitz C. Dyslipidemia and the risk of Alzheimer’s disease. Curr Atheroscler Rep.2013;15(3):307. doi:10.1007/s11883-012-0307-3

32. Scott RA, Scott LJ, Mägi R, et al. An Expanded Genome-Wide Association Study of Type 2 Diabetes in Europeans. Diabetes.2017;66(11):2888–2902. doi:10.2337/db16-1253

33. Shungin D, Winkler TW, Croteau-Chonka DC, et al. New genetic loci link adipose and insulin biology to body fat distribution. Nature. 2015;518(7538):187–196. doi:10.1038/nature14132

34. Sims R, van der Lee SJ, Naj AC, et al. Rare coding variants in PLCG2, ABI3, and TREM2 implicate microglial-mediated innate immunity in Alzheimer’s disease. Nat Genet. 2017;49(9):1373–1384. doi:10.1038/ng.3916

35. Sparks DL. Cholesterol metabolism and brain amyloidosis: evidence for a role of copper in the clearance of Abeta through the liver. Curr Alzheimer Res.2007;4(2):165–169.

36. Staffaroni AM, Elahi FM, McDermott D, et al. Neuroimaging in Dementia. Semin Neurol.2017;37(5):510–537. doi:10.1055/s-0037-1608808

37. Stearns FW. One hundred years of pleiotropy: a retrospective. Genetics.2010;186(3):767–773. doi:10.1534/genetics.110.122549

38. Steele NZR, Carr JS, Bonham LW, et al. Fine-mapping of the human leukocyte antigen locus as a risk factor for Alzheimer disease: A case-control study. PLoS Med.2017;14(3):e1002272. doi:10.1371/journal.pmed.1002272

39. Warde-Farley D, Donaldson SL, Comes O, et al.The GeneMANIA prediction server: biological network integration for gene prioritization and predicting gene function. Nucleic Acids Res.2010;38:W214–220. doi:10.1093/nar/gkq537

40. Watanabe K, Taskesen E, Bochoven A, Posthuma D. Functional mapping and annotation of genetic associations with FUMA. Nature Communications.2017;8(1):1826.doi:10.1038/s41467-017-01261-5

41. Westra H-J, Peters MJ, Esko T, et al. Systematic identification of trans eQTLs as putative drivers of known disease associations. Nature Genetics.2013;45(10):1238–1243. doi:10.1038/ng.2756

42. Willer CJ, Schmidt EM, Sengupta S, et al. Discovery and refinement of loci associated with lipid levels. Nat Genet.2013;45(11):1274–1283. doi:10.1038/ng.2797

43. Yokoyama JS, Karch CM, Fan CC, et al. Shared genetic risk between corticobasal degeneration, progressive supranuclear palsy, and frontotemporal dementia. Acta Neuropathol.2017;133(5):825–837. doi:10.1007/s00401-017-1693-y

44. Yokoyama JS, Wang Y, Schork AJ, et al. Association Between Genetic Traits for Immune-Mediated Diseases and Alzheimer Disease. JAMA Neurol.2016;73(6):691–697. doi:10.1001/jamaneurol.2016.0150

45. Yu J, Vodyanik MA, Smuga-Otto K, et al.Induced pluripotent stem cell lines derived from human somatic cells. Science.2007;318(5858):1917–1920. doi:10.1126/science.1151526

46. Zhang Y, Sloan SA, Clarke LE, et al. Purification and characterization of progenitor and mature human astrocytes reveals transcriptional and functional differences with mouse. Neuron.2016;89(1):37–53. doi:10.1016/j.neuron.2015.11.013

